# Multi omics aided small RNA profiling of wheat rhizosphere and their potential targets in contrasting soils for *Rhizoctonia solani*-AG8 suppression

**DOI:** 10.1101/2024.06.17.599338

**Authors:** Roshan Regmi, Shivangi Panchal, Marcus Hicks, Stasia Kroker, Jonathan Anderson, Gupta Vadakattu

## Abstract

Next-generation sequencing helps describe microbial communities in rhizosphere environments, but understanding rhizosphere-plant interactions’ synergistic effects on plant traits and health outcomes remains challenging. This study analyses rhizosphere sRNAs’ ability to manipulate host gene targets in plants grown in suppressive (SP) and non-suppressive (NSP) soils with an integrated multi omics dataset. The results showed that rhizosphere sRNAs exhibited specific compositional features that may be important for rhizosphere-plant interaction. Small RNAs, less than 30 nt in size, were predominant in both samples, with a 5-prime bias towards cytosine enrichment, suggesting potential association with wheat specific argonauts. Mapping of sRNA reads to microbial metagenomes assembled draft genomes from SP and NSP soils showed sRNA loci were differentially expressed (DE) between the soils with contrasting disease suppressive capacities. In total, 96 and 132 non redundant rhizosphere sRNAs were abundant in SP and NSP rhizosphere communities, respectively. While 55 known bacterial sRNA loci were predicted from both SP and NSP metagenomes, 127 sRNAs originated from these loci were differentially expressed. Global wheat target prediction and functional analysis from DE rhizosphere sRNAs showed both soil type specific and common pathways. Upregulated NSP sRNAs target metabolic pathways, secondary metabolite biosynthesis, MAPK signalling, while SP sRNAs target glycerophospholipid metabolism, pathways such as polycomb repressive complex, starch/sucrose metabolism, and plant-pathogen interactions were targeted by both sets of sRNAs. This is the first study showing evidence for rhizosphere sRNAs and their corresponding plant transcripts in the context of biological disease suppression in agricultural soils.

**Importance:** Small RNAs (sRNAs) have gained attention in host-microbe interactions due to their diverse roles in controlling biological processes. Studies have identified numerous sRNAs with novel functions across various organisms. Echoing growing evidence of sRNAs in different plant-microbe interaction, we show an evidence of rhizosphere sRNAs regulating wheat genes in soil disease suppression context. This understanding could significantly enhance our comprehension of gene regulation in biological functions, potentially paving the way for the development of microbiome-based methods to influence host traits. Understanding the microbiome community’s mechanisms in different environments offers opportunities to modify them for agriculture, including modifying farming practices, host genetics/immunity, and synthetic communities for disease suppression.

## Introduction

The rhizosphere is a highly complex and dynamic microhabitat, consisting of plant roots, soil and an integrated consortium of bacteria, fungi, eukaryotes, and archaea [1, 2]. The nature of interactions among them (mutualistic or antagonistic) is a driver of plant health and determines the functions and ecosystem services provided by rhizosphere microbiome communities [3]. Plants rely on the rhizosphere microbiome for specific traits and functions, while excretion of root exudates from plants provides a rich carbon source for the microbiome communities and influences their functional activity and diversity [4]. A wealth of knowledge is currently available on the taxonomy of microbial communities under different microbiome-plant interaction conditions, with a focus on bacteria and fungi community [4, 5], but not limited to, pathogens [6–8], symbiont rhizobia [9, 10] and mycorrhizal fungi [11]. In addition, microbiome profiling of the rhizosphere under different conditions such as nutrients availability [12], host genotype variation [13, 14] and stress [15, 16] also of interest. However, detailed functional mechanisms controlling such interactions are still obscure. With the advent of next generation sequencing, it is now possible to understand the microbiome continuum present in the complex rhizosphere environment and the nature of their communications that might impact on the functional capability of the whole system.

Metagenomics and meta-transcriptomics are useful tools to understand the functional potential and transient responses of microbial communities. However, neither approach provides a complete picture of gene regulation and inter-species interactions as both prokaryotes and eukaryotes have been shown to utilise sRNA mediated post transcriptional gene regulation to control biological process including development and growth, maintenance of genome integrity and response to abiotic and biotic stress [17]. Regulation by sRNAs is particularly evident in processes that require rapid responses to external signals that are present within host environments e.g., envelop stress response, carbon, and metal ion limitation [18, 19]. sRNAs are epigenetic regulators with a size of 18-30 nt in eukaryotes and 50-500 nt in prokaryotes that regulate the target genes through sequence complementarity. Regardless of their biogenesis mechanism, both prokaryotic and eukaryotic sRNAs regulate important biological activities [20]. Given our understanding on the widespread involvement of sRNAs during plant-microbe interaction [21, 22], it is suggested that sRNA mediate the modulation of host metabolic pathways and defense responses in rhizosphere microbiome/plant interactions [22].

To date, biochemical root exudates and signal molecules have been identified as central regulators of rhizosphere microbiome assembly and function, but the role of sRNAs in the formation of such assemblages and their functions has not been investigated [20]. For example, knowing how sRNAs secreted by bacteria and roots control interactions between plants and the belowground microbiome could help decipher the molecular mechanism of rhizosphere-plant interactions.

Commonly known as cross-kingdom RNA interference mechanism (cross-kingdom RNAi), sRNA-mediated inter-species communication has been reported for some microbe-plant interactions [21] such as modulation of plant immunity genes and pathogen virulence-related genes through pathogen and plant sRNAs, respectively [23, 24]. For example, the necrotrophic fungus *Botrytis cinerea* secreted sRNA effectors hijack the Arabidopsis immune system by suppressing several key plant defense genes. Meanwhile, Arabidopsis sRNAs silenced B. cinerea virulence genes through host-induced gene silencing [24, 25]. Similarly, *Fusarium graminearum* influenced the wheat immune system by silencing resistance-related genes encoding the “Chitin Elicitor Binding Protein” [26]. Furthermore, silencing of one or two pathogen effectors through RNAi can induce quantitative resistance in wheat during *Blumeria graminis* f.sp. *tritici* infection [27]. All of these studies are focused on the above ground fungi-host plant interactions. Recently, evidence on rhizobia sRNA modulation of the host through sRNA trafficking has been reported, an example for bacterial sRNA-mediated mechanism for prokaryotic-eukaryotic interactions [28]. *Trichoderma*, a beneficial fungus, has been shown to suppress the effects of soil-borne fungal pathogens and enhances plant immunity through the transmission of its sRNA [29]. However, a comprehensive investigation of the role of rhizosphere sRNAs in plant-microbe interactions including their potential for a role in disease suppression remains lacking [30]. Soilborne disease suppression is one of the key soil functions where the soil/rhizosphere/endophytes microbiome suppresses the effect of the fungal pathogens even in the presence of the susceptible host [6, 7, 31, 32]. The identification and detailed study of disease-suppressive soils can help to discover beneficial microorganisms with new antimicrobial and plant-promoting properties and help identify new farming practices with reduced disease impacts. In this study, we used a Rhizoctonia-wheat microbiome system in soils differing in their ability to suppress rhizoctonia root rot disease caused by *Rhizoctonia solani* AG-8 to study the role of sRNAs rhizosphere interactions. Rhizoctonia root rot is devastating soil borne disease that causes 77 million annual losses to wheat and barley production in Australia and a major yield limitation to cereal crops in US Pacific Northwest [33]. Currently, as there are no disease resistant wheat varieties or reliable chemical control options, better understanding of disease suppressive activity is important for sustainable management of this disease [34, 35]

The main objective of this study was to understand the differences in rhizosphere sRNAs between soils with different disease suppression activities and examine the functions and expression of plant genes targeted by these sRNAs. For this, sRNA sequences from wheat rhizosphere samples in disease suppressive and non-disease-suppressive soils along with shotgun metagenomics data for predicting sRNA loci and wheat root transcriptomics for sRNA target predictions was generated.

### Microcosm set up and nucleic acid extraction

Topsoils (0-10 cm) were collected from farmer fields at Avon and Poochera (Eyre Peninsula) in South Australia. Soil in the Avon field (suppressive) has a history of disease suppressive abilities for soilborne diseases such as rhizoctonia [31] while Poochera field soil (non-suppressive) has no significant disease suppression ability. Their respective disease suppressive abilities were further confirmed in controlled environment pot bioassays through Rhizoctonia root rot based on disease trait analysis. We refer to SP for the suppressive Avon soil and NSP for Poochera non-suppressive soil from here onwards throughout the manuscript. One kilogram of soil was packed into 1.5 L pots and inoculated with one week old (one cm) mycelial plugs (at 1 cm depth) grown on ½ Potato Dextrose Agar and incubated for a week at 15 ℃. The pots were watered twice a week to keep the moisture level at 12% w/w for SP and 16% w/w for NSP soil. After 7 d incubation, 4 seeds per pot were sown and the soil surface covered with polyethylene beads to reduce quick evaporative loss of water. Three seedlings per pot were retained after germination. After 7 weeks, rhizosphere and root samples were collected for nucleic acid extraction. Briefly, the plants removed from the pots were shaken gently to remove the bulk soil. Rhizosphere soils (closely attached to roots) and roots were separated by gently shaking 3-4 times, roots separated from the rhizosphere soils were washed briefly by a sterile water. Both rhizosphere soils and roots were immediately frozen in liquid nitrogen and stored at −80 ℃ until nucleic acid extraction. To maximize the coverage of sequences/transcripts, plants from six pots were pooled together and treated as a single biological replicate. Co-isolation of rhizosphere DNA/RNA was done using RNeasy PowerSOIL and DNA (deoxyribonucleic acid) elution kits (Qiagen) from three separate biological replicates with RNA extraction from corresponding root samples using RNeasy Plant Mini Kit (Qiagen). The quantity and RNA integrity of rhizosphere RNAs was checked using qubit and running in gel electrophoresis and tape station. RNA from three individual rhizosphere biological replicates from SP and NSP soils and their corresponding rhizosphere and root samples were sent to Australian Genome Research Facility (AGRF) for small RNA and plant RNA sequencing, respectively. The sRNA libraries involved Nextflex v3 small RNA workflow and sequenced in single end read mode (1 x 100 bp) while RNA sequencing was conducted on a NovaSeq in paired end mode (2 x 150 bp) after plant ribosomal depletion. DNA from three same biological replicates used for sRNAs were pooled together and sent to Phase Genomics USA (www.phasegenomics.com) for shotgun metagenome sequencing. Metagenomics library was prepared using Proximeta library preparation reagents from the Phase Genomic ProxiMeta Kit v4.0 and was sequenced on a NovaSeq in a paired end mode (2 x 150 bp) to a sequencing depth of 100 million reads. In summary, six rhizosphere sRNA libraries with their corresponding root RNA libraries and two shotgun metagenomics libraries one for each SP and NSP soil were developed in this study.

### Bioinformatics analysis

The quality of raw reads was assessed with fastqc tools [36] and trimmed/quality filtered to remove adaptors and low-quality bases with a bbmap.sh script from bbmap program (version 39.01) [37]. Furthermore, rRNA reads were removed using reference sequences from a SILVA database [38]. For sRNA cluster identification, clean sRNA reads were mapped to references using ShortStack (version 3.8.5) program [39]. ShortStack first identified significant target coverage based on target depth. Major alignments were then extended upstream and downstream. Overlapping areas were then combined into clusters. The most abundant sRNAs at each cluster were called major RNAs. These major RNAs were treated as putative sRNA sequences. Root RNAs were *de novo* assembled using Trinity software (version 2.13.2) [40]. Six individual libraries were assembled, and the assembled genome in fasta file was used as a reference to map back the individual library to find differentially expression genes. The count table of mapped reads to each assembled transcripts were produced using RSEM (RNA-Seq by Expectation-Maximization) available in the trinity software package. Biological replicates were compared for the SP and NSP libraries with Pearson correlation matrix calculated in Trinity based on transcript expression values. RSEM outputs two files for isoform-level and gene-level estimates, providing abundance estimates in terms of non-integer values. The first approximation estimates the number of fragments derived from a given isoform or gene, based on maximum likelihood abundances.

Shotgun reads were filtered and trimmed for quality and then individual or co-assembled with MEGAHIT (version 1.2.9) [41] with the k-mer range set to 21,41,61,81,88 to account for sample complexity “kmin-1 pass --presets meta-large --k-list 21,41,61,81,99 --no-mercy --min-count 2”. The assembled contigs were binned using MaxBin2 (version 2.2.7) [42]. At first coverage file was generated by mapping individual libraries back to the assembly using bbmap.sh and pileup.sh command from bbmap. Thus, generated abundance file was supplied to MaxBin2 with a command “-contig final contigs. fa -abun abundbace.txt” which produced the output folder of the assembled bins. All individual assembled bins were concatenated to get a single fasta file and used as a reference file for sRNA reads mapping. We have thus created three de novo assembled draft metagenomes (MAGs) each for SP, NSP and co-assembled bins. Known bacterial sRNAs were annotated from assembled SP and NSP metagenomes using infernal program and RFAM database [43] using the script from Regmi et al. 2022 [44]. The sRNA libraries were mapped back to annotated bacterial sRNA loci in metagenomes using bowtie2 [45]. The RFAM annotated sRNA loci were extracted from SP and NSP draft metagenome. The fasta file of each loci (common and specific to each metagenome sample) was parsed to Kraken2 [46] and run with a command “kraken2 -db $dbr --threads 12 --output sRNAlociKraken.txt --report kraken Output sRNAlocibp.fa”, while dbr represents all the sequences from Kraken2 database, sRNAlociKraken.txt was the output text file and sRNAloci.fa was an input file. The results from Kraken2 were visualized and raw reads were extracted using pavian platform [47]. psRNA target server [48] was used for in silico target prediction upregulated rhizosphere sRNAs in the wheat transcriptomics. This tool is based on specific pairing patterns between plant sRNAs and their targets at 10th and 11th nucleotide from 5’ region of complementary sRNAs. Based on the psRNA results, sRNA-target recognition, and the e-value was computed for each alignment and those with an e-value <= 5 were kept for functional enrichment analysis. It is to be noted that, this hypothesis is grounded based on bacterial sRNAs repress the plant target genes in the same manner, through sequence complementarity. It is possible there might be yet unknown mechanisms how these interactions might occur during prokaryotic-eukaryotic interaction. For 127 DE bacterial sRNAs, we could not use a psRNA target server as this program only accepts sRNAs with a size below 25 nt, used a IntraRNA for RNA pairing information which is optimized for a bacterial sRNA target recognition [49].

The identified target genes were subjected to gene ontology (GO) and KEGG (Kyoto Encyclopedia of Genes and Genomes) analysis using David functional annotation tools [50],where predicted targets were input as gene lists and all wheat Ensemble transcripts as background list. The resulting GO terms and KEGG of the target genes were downloaded and plotted in R (4.1.2) using ggplot2 package [51]. The enrichment of KEGG pathways was considered in terms of fold enrichment, p-values and the number of genes present in the pathway. The fold enrichment is the ratio of the number of target genes in the pathways and the number of all the genes annotated in the pathways. For de novo assembled DE targets, coding sequences of sRNA targets were annotated with eggNOG [52]. Those transcripts with identified pathways were manually inspected, and small subset was confirmed through real time quantitative polymerase reaction (RT-qPCR). Transcripts were chosen based on the target recognition sites observed in upregulated transcripts in SP trasncriptomes from both upregulated sRNAs and NSP sRNAs. In total, five references were used to find sRNA clusters from the rhizosphere sRNAs, among which three were developed in this study; draft MAGs from SP, NSP and co-assembled, wheat mace IWGC genome and *Rhizoctonia solani Ag-8* references were downloaded from Ensemble [53]. The schematic flowchart used for a bioinformatics analysis is shown in Figure 1.

**Figure 1.**
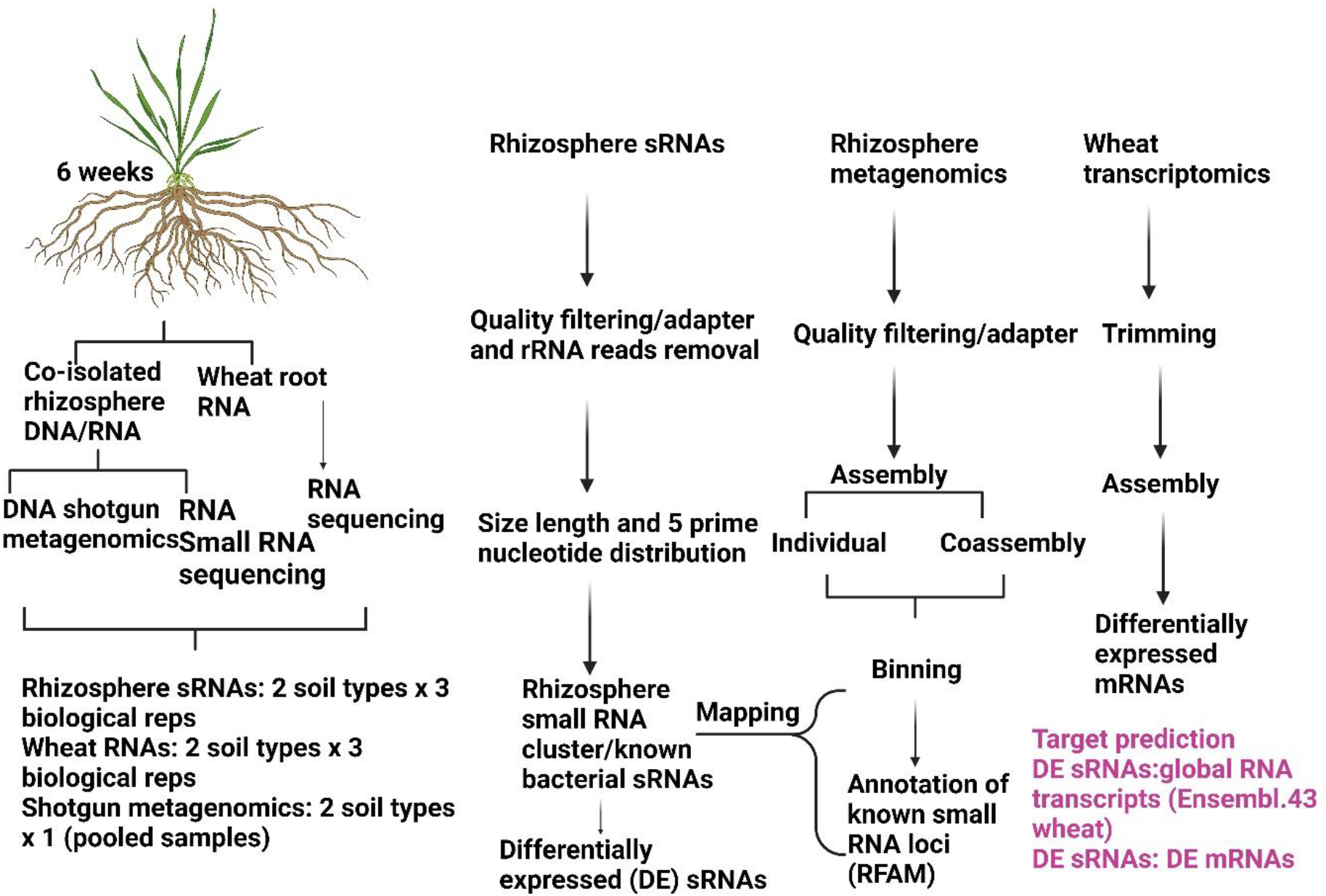
Data acquisition and bioinformatics pipelines used in this study.

### Real time quantitative polymerase chain reaction

Total RNAs were DNAse treated with DNAse enzyme (Invitrogen TM). One µg of DNAse treated RNAs were converted into cDNA using a protoscript first strand cDNA synthesis (New England Biolab TM). The resulting cDNA samples were subsequently diluted to 20 times before being subjected to RT-qPCR analysis. The reactions were run in 15 µl, with 1 µl of cDNA, 7.5 µl of Syber green master mix (BioradTM), 2 µl of forward and reverse primers, and 4.5 µl of nuclease free water. The reactions were run in CFX Opus Biorad 384 Real-Time PCR with an initial denaturation step at 95°C for 3 min followed by cycles of 95°C for 10 sec, 55°C for 10 sec, and 72°C for 30 sec. This cycle was repeated 39 times, and the procedure concluded with a last step at 65°C for 30 sec, followed by 95°C for 5 min for melting curve analysis. Each sample was analysed with three biological replicates and three technical replicates and repeated twice. The average value from two runs were taken for final relative expression analysis. The relative expression of the target genes was calculated using log (2-_ΔCt_) method with normalization to the wheat housekeeping actin gene. Primers used in this study are listed in Supplementary Table 1.

## Results

### Overview of rhizosphere small RNAs

To study the role of sRNAs in the interaction between plants and soil diseases, we analyzed sRNA sequences from two types of soils with different suppression abilities against the soilborne pathogen *Rhizoctonia solani* Ag 8 We generated 4.4 billion reads from six libraries and retained an average of 70% and 50% reads from SP and NSP samples, respectively. Length distribution and quality assessment were done for the sRNA dataset [54, 55] as random degradation of sRNAs yielded uniform size distribution, while specific biogenesis processes create size preferences [44]. For both SP and NSP samples, 23 nt and 24 nt were present in higher proportions than other size lengths (Figure 2) with 24 nt previously being reported to be characteristics feature of wheat and fungal sRNAs [56, 57]. Rhizosphere sRNAs are a mixture of both prokaryotic (bacterial) and eukaryotic (fungal and plant) molecules where eukaryotic sRNAs are reported to be < 30 nt and bacterial sRNAs >30 nt but sRNA fragments below 30 nt [58] were also assigned to be of a bacterial origin with high throughput sequencing. Therefore, we expect sRNA sizes of less than 30 nt predominantly belonged to a bacterial origin and is of a particular interest from rhizosphere microbiome perspective, as we are interested in knowing bacterial specific sRNAs and how they manipulate the host genes. Also, comparing these size length characteristics with a previous reported studies on sRNAs would help to provide an indication of the quality of our dataset. Size length analysis revealed diverse sRNA heterogeneity in the rhizosphere, highlighting bacterial-originated fragments. sRNA distribution featured significantly more sequences below 30 nt compared to those above 30 nt. Interestingly, there was size length variation for some sRNAs between two samples, such as 23-28 nt enrichment in SP samples and 29 nt predominance in NSP samples. Both samples showed an enrichment at 100 nt. Notably, most sRNAs displayed a cytosine bias at the 5’ ends. (Fig 2 insert).

**Figure 2.**
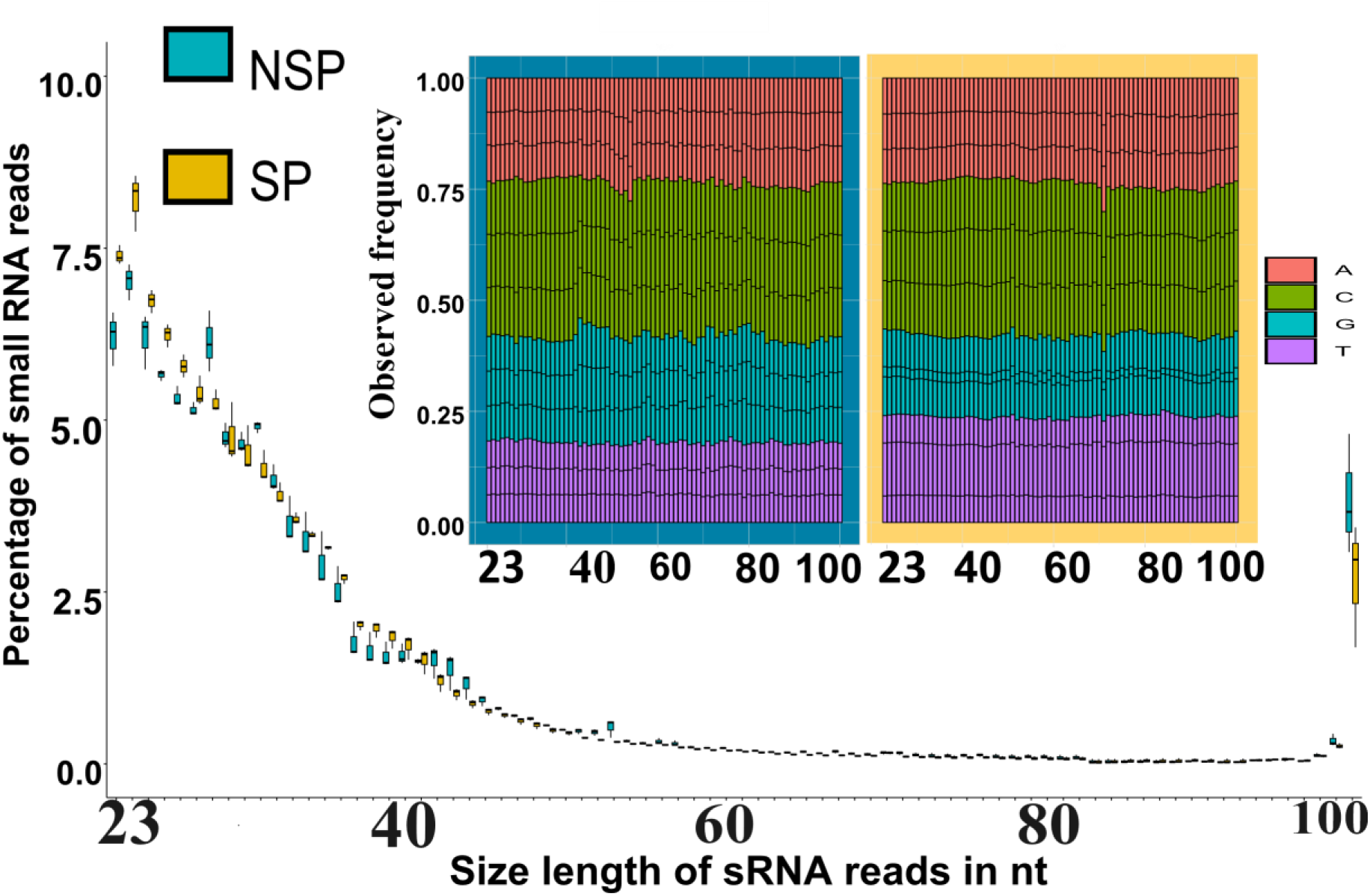
Rhizosphere small RNA characteristics analysed in terms of size length and 5 prime nucleotide bias from SP-suppressive and NSP-non suppressive sample. Box plot shows the first and third quartiles, and the centre line indicates the medians (n=3), and the whiskers extends to the most extreme data points. Figure insert represents five prime nucleotide bias of rhizosphere sRNAs, x-axis shows the size length of sRNAs from Fig 1A, and y-axis shows the percentage of A (pink), C (green), G (blue) and T (purple) at 5 prime positions for the given size length.

### Cluster based approach for rhizosphere small RNA identification

We mapped sRNA libraries to draft metagenomes assembled genomes to identify the potential origins of the sRNA clusters. A total of 2,698, 2,705, and 2,534 clusters were predicted from NSP, SP, and co-assembled bins. Among these, only 222 and 36 clusters were found in wheat and *Rhizoctonia* genomes, respectively. Two wheat sRNAs were differentially expressed between the SP and NSP libraries, while no differential expression was observed for sRNAs originating from RS. Differential expression analysis of bacterial derived sRNAs revealed 96 and 132 non-redundant sRNAs with higher abundance in SP and NSP MAGs, respectively (Figure 3A). Differentially expressed sRNAs between SP and NSP soils were utilized to predict downstream silencing targets in wheat.

**Figure 3.**
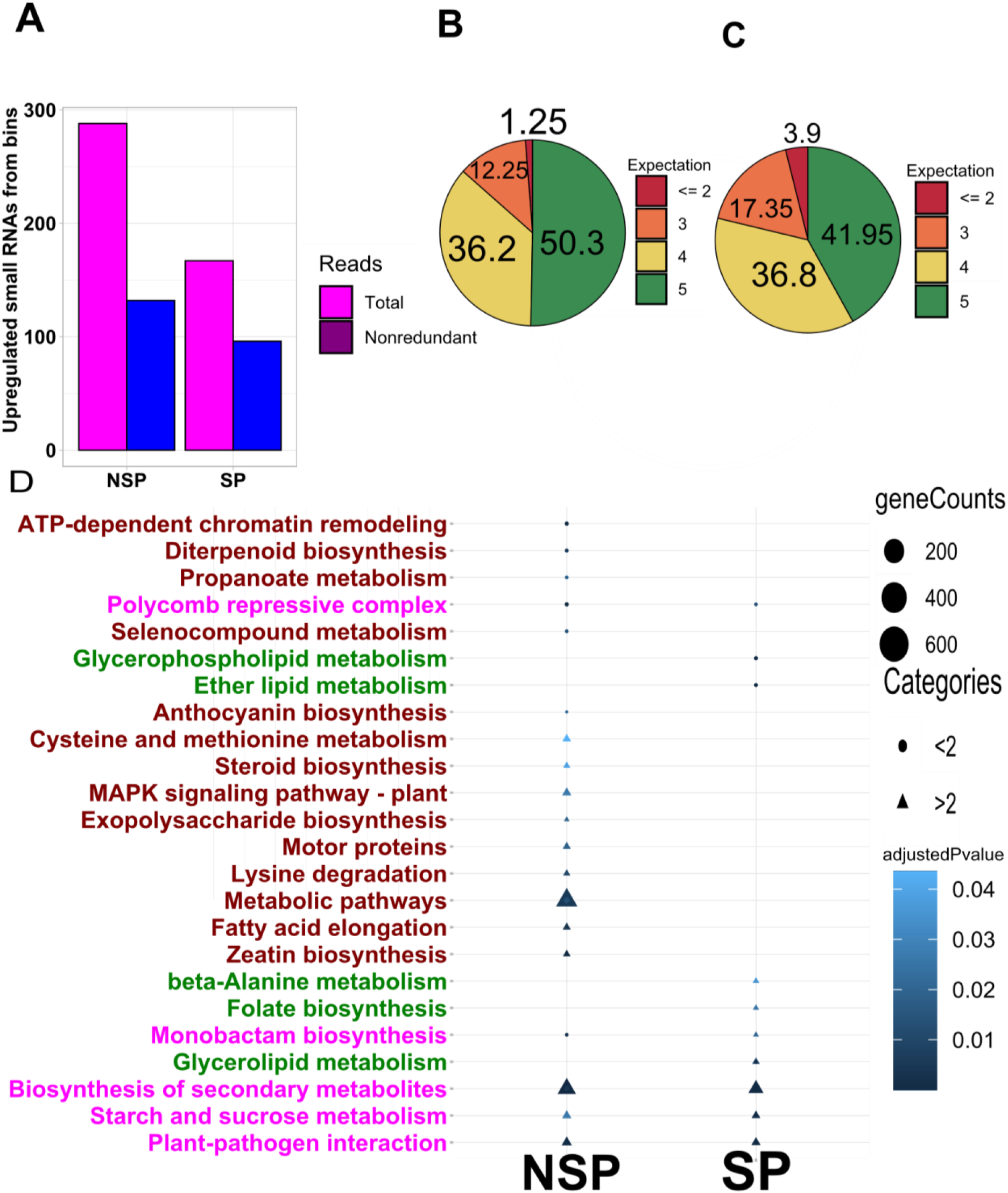
Differentially expressed rhizosphere small RNAs and their putative wheat targets. Number of total and unique upregulated sRNAs from SP-suppressive and NSP-non suppressive rhizosphere (A). Pie-chart showing the percentage of different categories of wheat targets predicted from psRNA target server based on expectation score from (B) upregulated SP sRNAs and (C) NSP sRNAs. KEGG pathway analysis of upregulated sRNAs potential wheat gene targets of differentially expressed sRNAs wheat genes with an expectation score of = < 2 and = > 2 (D). The x-axis shows fold enrichment, size of the circle denotes the gene counts find in given pathways. The terms in red are those only present in targets of upregulated NSP sRNAs, while the term with pink is present in targets of both sets. The terms that are present in SP targets only are shown in green. Circle represents NSP with an expect score less than 2, and triangle denotes SP an expect score between 2 and 5 targets.

### Global overview of rhizosphere sRNAs potentially targeting wheat genes

We predicted wheat target genes for differentially regulated bacterial sRNAs. A total of 12,873 target sites were predicted from both sets, with around half having an expectation score of 5, indicating weak sRNA:mRNA pairing (Figure 3B). Low expectation score targets are usually considered genuine sRNA targets [59]. Therefore, to ensure accuracy in our gene analysis, we focused on sRNA target sites with low (<= 2) and high (>=2) expectation scores separately. Pathway analysis was performed to estimate target gene functions. GO terms describe gene product functions, while KEGG classification provides molecular level information through gene and molecule interactions [60]. Nine KEGG pathways were found for NSP sRNA targets and four for SP sRNA targets. Both sets shared the “Polycomb repressive complex” pathway, while lipid metabolism was enriched in SP targets (Figure 3D). The KEGG pathway analysis of targets with an expectation score between 2 and 5 comprised MAPK signalling and metabolic pathways related genes from NSP targets while some terms like plant-pathogen interaction, sucrose and starch metabolism, secondary metabolite synthesis were present in both sets (Figure 3D).

### Mapping of RNA-seq to Ensemble plant wheat transcripts

Mapping of RNA-seq reads from roots to the reference wheat genome retrieved from Ensembl plant revealed only 30-35% of reads were mapped. This resulted in finding the upregulation of 1734 transcripts and the downregulation of nine transcripts in SP samples in comparison to NSP samples (Supplementary Table 2). There might be more altered expression but could not be analysed as all reads could not be mapped. The functional annotation of upregulated SP transcripts resulted into 14 significantly enriched biological process “ATP synthesis”, Hydrogen ion transport”, “Ion transport”, “Protein biosynthesis”, “Glutathione biosynthesis”, “Transport”, “Photosynthesis”, “mRNA splicing”, “Electron transport”, “mRNA processing”, “mRNA capping”, “Methionine biosynthesis”, “Ribosome biogenesis” and Asparagine biosynthesis”. In addition, five KEGG pathways were assigned to these transcripts, including “Ribosome”, “Oxidative phosphorylation”, “Spliceosome”, “Endocytosis” and “Alanine, asparate and glutamate metabolism”.

### Rhizosphere sRNAs and *de novo* host transcriptomics target pairing

Mapping RNA-seq data to closely related species or global transcriptomes may result in loss of species-specific gene information and might not accurately reflect the effect of gene expression on the phenotype or trait analysis. Moreover, one of challenges in transcript quantification from RNA-seq data is handling of reads that align to multiple genes or isoforms. This is even important for quantification with *de novo* transcriptome assemblies in the absence of species-specific sequenced genomes [61]. The accuracy of predicting sRNA targeting transcripts by sRNAs is enhanced when paired with bespoke target transcripts from the same experiment. Taking this into account, *de novo* transcript assembly was performed to improve mapping success, resulting in 100% reads from all libraries being mapped to a de novo transcriptome. The differential expression analysis of individual samples revealed 314 DE transcripts between SP and NSP samples, with 205 and 109 upregulated in SP and NSP samples, belonging to 51 and 31 longest isoforms, respectively (Figure 4A). We converted 314 DE nucleotide sequences into protein coding sequences to obtain functional information about DE transcripts, resulting in 85 and 8 in SP and NSP samples, respectively. Among 85 protein coding sequences in SP samples, 31 were annotated (Supplementary Table 3), while 54 did not shown any similarity to putative functions. The annotated transcripts related to various functions, such as “RNA mediated transposition”, “ATP synthesis”, “Glycotransferase” and “photosynthesis (chlorophyll)”. These functions were broadly categorized to cluster of orthologous groups of protein (COG) categories of “(C) Energy production and conversion”, “(J) Translational, ribosomal structure and biogenesis”, “(L) Replication, recombination and repair”, “(H) coenzyme transport and metabolism”, (O) Post translational modification, protein turn over, chaperons. The functions predicted to be upregulated in SP samples mostly corroborated to those identified from transcriptomics analysis using a reference from Ensemble Plants. The eight protein coding sequences with higher abundance in NSP samples did not show similarity to proteins with known functions. However, blasting the nucleotide sequences of the 109 transcripts with higher abundance in NSP samples in NCBI blast revealed homology mostly to repeat sequences in wheat genome. The psRNA target server was further used to determine if the upregulated small RNAs (sRNAs) were paired with differentially expressed wheat transcripts. The sRNA:mRNA pairing ratio was lower in NSP samples compared to SP samples, indicating that NSP sRNAs regulate more targets per sRNA (Figure 4B). The lower ratio of sRNA:mRNA in NSP samples might be due to repeat elements. Five transcripts with common functions from *de novo* and global transcriptomics were randomly selected for validation through RT-qPCR, and three of them were found to be significantly upregulated in SP samples and provides similar expression patterns from RNA-seq data (Figure 4C).

**Figure 4.**
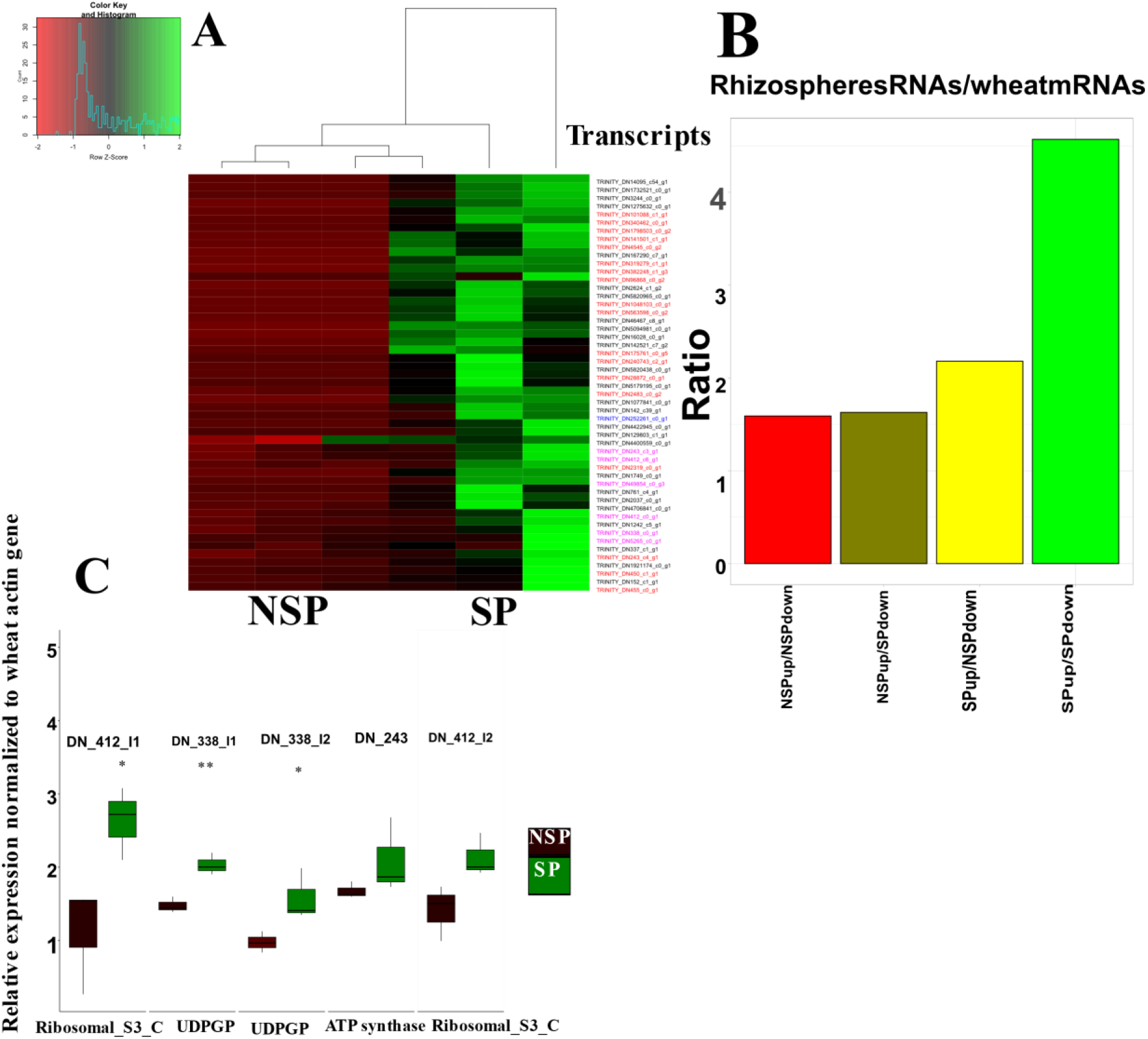
Host transcriptomics analysis and qPCR validation. (A) Heat map of upregulated 51 transcripts upregulated in suppressive soils identified. (B) sRNA:mRNA pairs of upregulated sRNAs and mRNAs showing the ration of number of targets regulated by each unit of sRNAs. (B) NSP up/NSP down indicates the average number of wheat transcripts that are downregulated in the NSP samples for each corresponding rhizosphere sRNA observed in the NSP samples. SPup/Spdown indicates the average number of wheat transcripts that are downregulated in the SP samples for each corresponding rhizosphere sRNA observed in the SP samples. qPCR validation of transcripts combinedly target by upregulated sRNAs from both sets. Box plots indicate and whiskers extend to the most extreme data points (C).

### Identification of known bacterial sRNAs from rhizosphere sRNA sequences and metagenomes

After the identification of rhizosphere sRNAs and their potential targets in wheat, we hypothesised that some sRNAs might belong to characterised bacterial sRNA families and potentially differ in abundance between SP and NSP samples. To this end, we first annotated known bacterial sRNAs in our MAGs using the RFAM database. The sRNA read libraries were subsequently mapped to the metagenome contigs containing the characterised sRNAs to assess their expression in SP and NSP samples. The RFAM [62] database contains 3701 bacterial sRNA sequences originating from 3215 bacterial species. Metagenome contigs from SP samples contained 60 known sRNAs whereas NSP contigs contained 67 sRNAs (Figure 5A). Among these sRNA loci, 55 were common in both samples, while 12 and 5 sRNAs were present only in NSP and SP MAGs, respectively. Most of the annotated sRNAs common to both NSP and SP samples belonged to the families of toxic beta proteobacteria sRNAs and 5_ureB. Toxic beta proteobacteria sRNAs are a family of trans-encoded sRNAs exclusively found in intergenic regions of betaproteobacteria while 5_ureB targets are found in the urease gene family responsible for nitrogen cycling within soils [63, 64]. We classified the 55 common sRNA loci contigs using a Kraken2 database, primarily originating from proteobacteria. Sixteen sequences were identified up to species level (Figure 5B), including *Amycolatopsis sp. AA4*, *Pseudorhodoplanes sinusperici*, *Paracoccus sanguinis*, *Stappia indica*, *Tabrizicola piscis*, *Janthinobacterium agaricidamnosum*, *Massillia abdiflava*, *Varivorax paradoxus*, *Ocanisphaera profuna*, and six uncultured prokaryotes. Similarly, among five sRNA loci specific to SP samples, three were classified up to a species level which include “*Rhizobium sp. NXC24*, *Massilia putida*, and one uncultured prokaryote. For NSP specific sRNA loci, eight were classified up to a species level, which contain *Amycolatopsis sp. AA4*, *Streptacidiphilus* sp. DSM 106435, *Streptomyces violaceoruber*, *Devosia* sp. A16, *Rhizobacter gummiphilus* and three uncultured prokaryotes.

**Figure 5.**
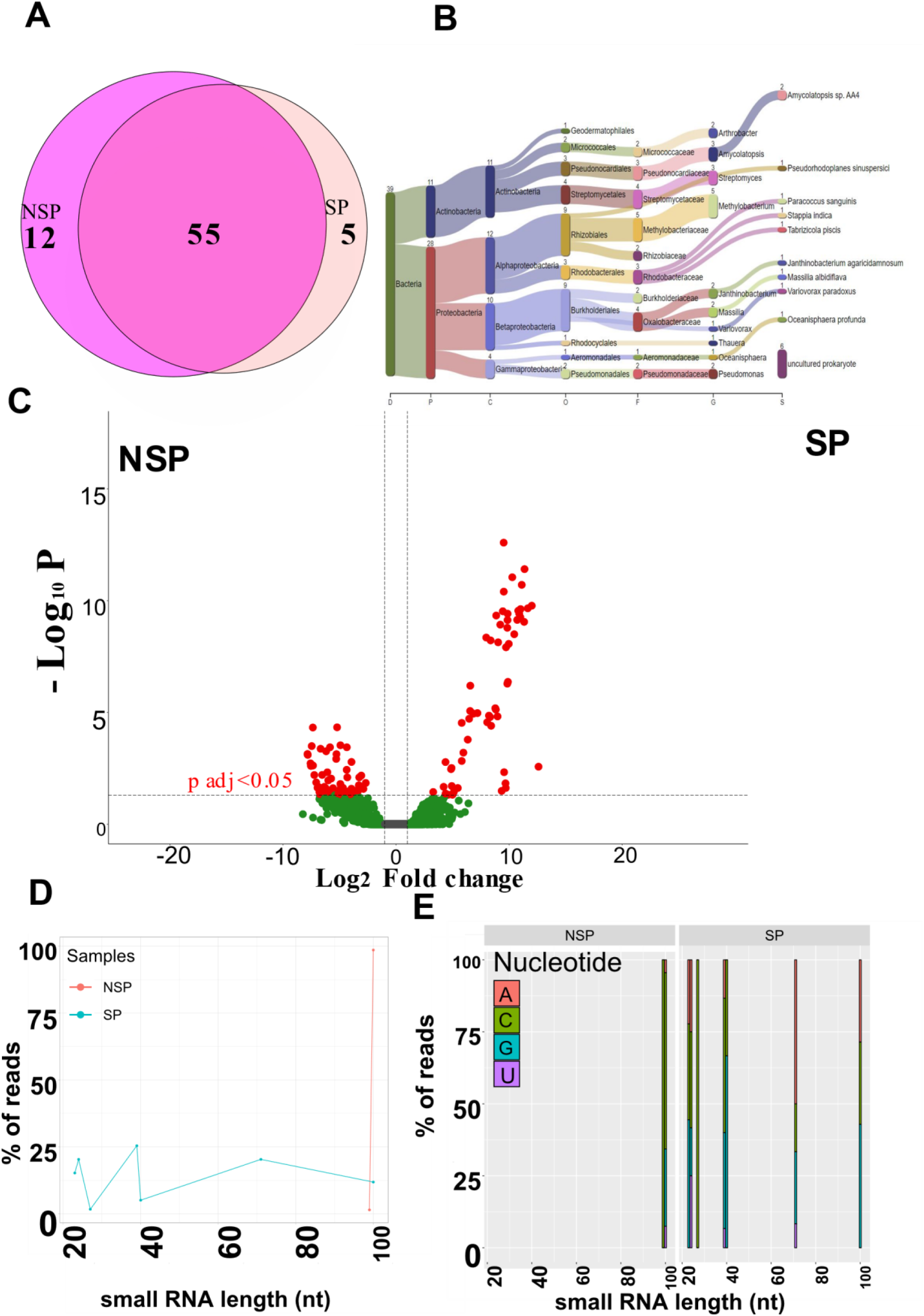
Identification of known bacterial sRNAs loci from assembled metagenomes (MAGs). Venn diagram showing the identification of bacterial sRNA loci from RFAM database in suppressive and non-suppressive draft metagenomes. (A). Classification of 55 common sRNA loci using Kraken2 database, among 55, 45 sequences were classified among which 39 were classified to bacterial sequences (B). Volcano plot showing the differentially expressed sRNAs mapped to the MAG (metagenome assembled genomes) bacterial sRNAs loci, log2 fold change plotted as a function of the negative logarithm of P value (C). The red dots indicate the significantly differed sRNAs with log2 fold change of <1 and >1 and P adj value of < 0.05. The green dots show sRNAs that were only significant in terms of log2 fold change not of P adj value < 0.05. The grey dots represent sRNAs present in both SP and NSP samples (C). Size length distribution of 59 and 68 upregulated sRNAs in SP and NSP, respectively (D). 5 prime nucleotide bias of sRNAs from (D).

Bacterial sRNAs could be produced from their precursor longer fragments of dsRNAs [65, 66]. To investigate if any of the 55 characterised sRNA loci that were common to both SP and NSP samples showed any differential expression, we mapped our sRNA libraries to the metagenome contigs containing the 55 annotated bacterial sRNAs loci. As a result,18176 sRNA reads from six sRNAs libraries were mapped, with 59 and 68 sRNAs significantly enriched in SP and NSP samples, respectively (Figure 5C). To understand the features of these DE sRNAs, we investigated their size length and 5 prime nucleotide characteristics. Upregulated SP sRNAs had more diversity of size lengths compared to NSP upregulated sRNAs (Figure 5D). Additionally, the 5 prime nucleotide composition of sRNAs varied between the two types of soils suggesting different sRNA processing mechanisms are at play (Figure 5 E).

### Prediction of the secondary structure of bacterial small RNA loci

The study of RNA secondary structure is crucial for comprehending the functions and regulation of RNA transcripts, as it provides insight into their folding patterns and potential ligand-binding interactions [67]. Furthermore, bacterial sRNAs with lower folding energy are generally considered more functionally relevant [68]. Figure 6 shows the predicted sRNAs structures for five of the characterised sRNAs specific to the SP samples have highly conserved structures and more than two loops. sRNAs specific to the NSP and common also exhibited similar patterns.

**Figure 6.**
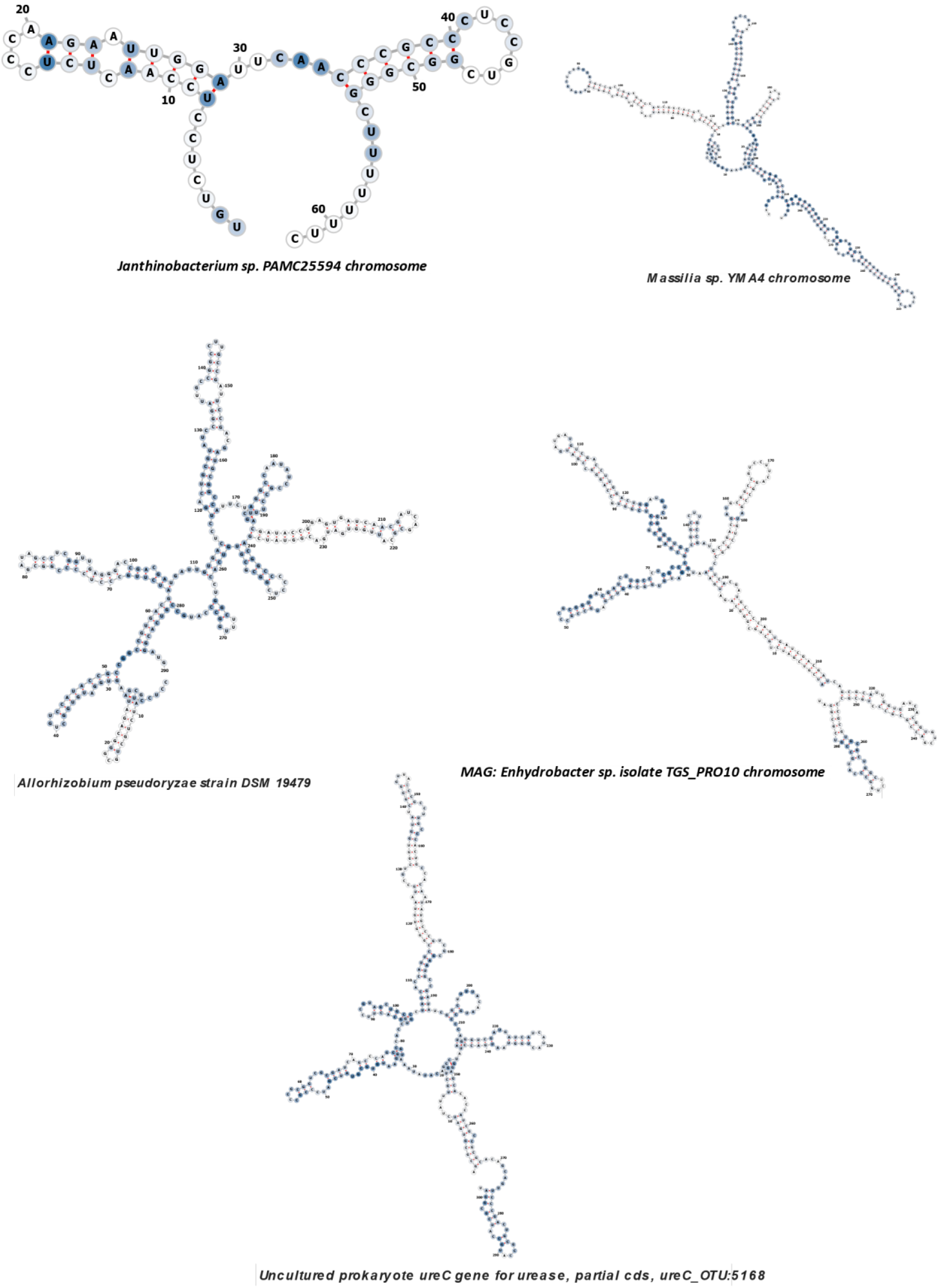
Secondary RNA structure of bacterial sRNA loci predicted only from SP-suppressive draft metagenome. The secondary structure is coloured according to the base-pairing.

## Materials and methods

### Discussion

Studies of microbiome mediated suppression of soilborne disease have until now been mainly focussed on describing differences in microbial composition including the research related to the suppression of rhizoctonia root rot disease [4, 7, 8, 56, 69, 70]. This study presents a new approach to gain insights into the regulation of rhizosphere-microbiome-plant interactions using a combination of rhizosphere sRNAs, metagenomics, and host transcriptomics. Results show that rhizosphere sRNAs were enriched with sRNA sizes below 30 nt in both samples along with a peak at 100 nt. Since the rhizosphere is a pool of both bacterial and eukaryotic sRNAs of related size, it has been considered difficult to provide a higher resolution of their origin [71], however metagenomics aided identification, as used in this study, provided additional support for the bacterial origin of some of the sRNAs. The proportion of sRNA clusters identified from bacterial MAGs were higher than those identified from wheat and Rhizoctonia, suggesting the importance of bacterial-derived sRNAs in these rhizosphere samples. In addition to these, rhizosphere soils would also have sRNAs from protozoa that might be involved in important functions [72].

Some size lengths were differentially distributed in two soil types, remarkably, 29 nt sRNAs were enriched in the NSP samples. Previously, 29 nt sRNAs derived from transfer RNA (transfer RNA-derived sRNA, tsRNA) of *Phytopthora pararasitica* were observed during infection of Arabidopsis roots [73] and have been implicated in the regulation of gene expression by inducing sequence-specific degradation of target RNAs. The relative abundance of 29 nt in NSP suggest these may be functional during pathogen infection and might be derived from Rhizoctonia tRNA fragments as pathogen infection was higher in the non-suppressive samples. Alternatively, these 29 nt sRNA may be derived from the wheat host as sRNAs derived from wheat tRNA were also reported to be induced during infection with the fungal pathogen *Fusarium graminearum* [74]. Further experimentation would be required to determine the origin and function of the 29 nt sRNAs in Rhizoctonia infection of wheat.

In eukaryotes, sRNAs are loaded into argonaute complexes to facilitate target gene silencing having a bias towards specific 5’ nucleotides [75]. Prokaryotic sRNAs differ from eukaryotic in their biogenesis but are mechanistically similar in suppressing targets. Recent discovery suggests that plant argonaute molecules such as argonaute5 are involved in processing bacterial-derived sRNAs, with corresponding regulation of transcriptional responses during symbiosis between Rhizobia and *Phaseolus vulgari*s [76]. Furthermore, sRNAs derived from rhizobial tRNAs were shown to be loaded into argonaute1 to control the nodulation process in legumes [28].

Understanding the 5’ nucleotide bias of bacterial sRNAs with corresponding argonaute molecules in eukaryotic hosts is important for understanding RNAi-based communication between them. Several argonaute genes have been identified in wheat [77], some of which are yet to be confirmed for their biological roles, with argonaute5 demonstrated to be involved in responses to aphid invasion [78]. Our data suggested rhizosphere sRNAs were biased towards cytosine in the 5’ nucleotide, suggesting argonaute5 in wheat may play a role in cross-kingdom RNAi between rhizosphere sRNAs and wheat transcripts. It is possible that rhizosphere-derived sRNAs with cytosine at 5’ end can hijack wheat transcripts via argonaute1/5. Further research on the role of argonaut5 using immunoprecipitation [79] from wheat will provide more insights.

Small RNAs regulate biological processes in prokaryotes and eukaryotes, possibly functioning in plant-microbe interactions through prokaryote-eukaryote cross talk [28]. Our *in silico* prediction identified potential plant targets of rhizosphere sRNAs, based on the hypothesis of prokaryotic-eukaryotic sRNA regulation. Initial findings showed plant stress and immune response pathways are regulated from rhizosphere/bacterial sRNAs. sRNAs often overlap with intergenic regions and with genomic regions, as they evolve from mRNA, random transcription, and tRNAs [58]. Our analysis of known bacterial sRNAs in our samples identified intergenic toxic betaproteobacteria and 5 ureases bacterial sRNAs, as common to both suppressive and non-suppressive soils. These sRNAs play a crucial role in regulating carbohydrate and amino acid metabolism [63]. 5’ urease sRNAs, present in the urease gene cluster and antisense to *ureB*, are essential for bacterial cell maintenance in tissues and colonization of host organisms [80, 81]. Urease catalyses the hydrolysis of urea into ammonia and carbamic acid [82]. It is possible that rhizosphere microbes could use urease to metabolise amino acids derived from plant root exudates in both healthy and disease plants. Additionally, these identified bacterial rhizosphere sRNAs also have a bias towards cytosine at the 5’ nucleotide, suggesting some may be transported to wheat and loaded into wheat argonaute5 to target wheat gene expression and bacterial outer membrane vesicles might conduit for sRNAs [83].

Interestingly, sRNA fragments mapped to a common 55 loci were differentially profiled in two soils. Under selection pressure, these sRNAs may be enriched in the rhizosphere and transported to the plant for fine tuning of target genes, as seen during root nodulation, where tRNA-derived bacterial sRNAs regulate the nodulation factor in legumes [28]

Further *in vivo* experiments involving plant targets and bacterial strains with sRNA sources may address this question. The Ribonuclease III (RNase III) family of endoribonucleases regulate gene expression by processing double-stranded RNAs. These are found in all kingdoms, where they play a major role in ribosomal RNA processing, post-transcriptional gene expression control, and viral defense [66]. Different fragment sizes of sRNAs in the two soils suggest different sRNA biogenesis strategies may be occurring in the two soils studied.

This study analysed the possibility of rhizosphere microbial sRNAs silencing of wheat genes by rhizosphere sRNAs through functional enrichment and pathways analysis. Both common and unique pathways were found in target genes of DE sRNAs, with the “lipid metabolism” KEGG pathway enriched in targets of sRNAs up regulated in SP samples. Lipids play a crucial role in plant defense by regulating signal transduction through jasmonate signalling, ROS metabolism, stomata closure, interacting with phytohormones and as precursors for bioactive defense molecules [84]. Interestingly, the upregulated NSP sRNAs target specific signalling pathways, such as Mitogen-activated protein kinase (MAPK-plant) and secondary metabolite biosynthesis pathways suggesting the silencing of plant immune response genes is higher in NSP samples as compared to SP samples. MAPK cascades are essential signalling modules in all eukaryotes, triggering cell division, differentiation, programmed cell death, and adaption to stress. In plants, these cascades are triggered by various stress stimuli and play crucial roles in hormonal and developmental signalling [85].

“Polycomb repressive complex”, “sugar and starch metabolism” and “plant-pathogen” interactions were assigned to targets of both set of sRNAs. Overall, the functions of target genes are associated with various functions, remarkably, some of the pathways associated with sRNAs with lower abundance in the SP samples (higher in NSP samples) were related to plant defence against pathogen infection [86–89], suggesting the potential for these wheat genes to be used as host markers for soil disease suppression. The expression of sRNAs are negatively correlated with their downstream targets [90]. Having lower SP sRNAs in SP soils and a higher number of upregulated genes provides an insight into the possibility of rhizosphere sRNA:mRNA regulatory involvement in disease suppression context.

In addition to *in silico* psRNA target prediction of DE sRNAs in a global wheat transcriptome, we also analysed transcriptomics of wheat roots generated in this study to understand how plant response in terms of their gene expression pathways while growing in SP and NSP samples. Wheat transcriptomics studies exhibit differential regulation of various gene pathways under different conditions [91–93]. Similarly in this study, wheat root gene expression differed between NSP and SP samples with a higher number of genes upregulated in SP samples. The gene pathways reveal differences in “oxidative phosphorylation”, “ribosome”, “spliceosome” and “endocytosis” between two samples. Endocytosis is a crucial process in plants that regulates plasma membrane dynamics, enabling them to respond to environmental cues and defend against pathogens [94]. Oxidative phosphorylation’s ability to precisely maintain metabolic homeostasis is crucial for the development of higher plants and animals [95]. Ribosomes and spliceosome are important pathways that are related to the regulation of transcription. The differential regulation of the ribosomal association of mRNA transcripts in an Arabidopsis mutant defect has been related to the wound response dependent on jasmonate [96]. Taken together, wheat grown in SP has ability to diversify transcriptome reprograming, potentially enabling plant adaptation to rhizoctonia infection, but further research is needed to determine its biological relevance.

Mapping of RNA-seq data to a closely related species or global transcriptomes might lead to a loss of information regarding species specific genes and incomplete information on the gene expression impact on the phenotype. Therefore, we conducted *de novo* assembly of root transcriptome data and used DE rhizosphere sRNAs to find their target sites on DE transcripts resulted from assembled transcriptomes. Some of the pathways that were upregulated in SP samples while aligning to the global transcriptome were also identified in *de novo* assembled transcripts being targets of sRNAs such as ATPase, ribsomal-s3 and chlorophyll related genes. Precisely, mapping of sRNA libraries to *de novo* assembled wheat transcripts, revealed SP samples had sRNAs targeting gene functions such as Glycotransferase, Chlorophyl related genes (Cytochrome oxidase), Aluminium malate transporter and ATP synthase. Pathogen induced plant Glycotransferase-like proteins have been shown to be associated with defence responses in plant [97]. Cytochrome oxidase genes were upregulated in disease resistant varieties in comparision to susceptible tomato varieties [98]. Moreover, cytochrome oxidase (chlorophyll) related genes were found to be downregulated and targeted by *Zymoseptoria* during infection of wheat [57]. Aluminium activated malate transporter was first discovered from wheat root tips, which was found to be involved in aluminium resistance by malate exudation in the soil [99]. Additionally, recent research has demonstrated that ALMT proteins can play important chemical roles in GABA (gamma-aminobutyric acid) signalling in plants [100]

Increased plant defense gene expression is a crucial early response to infection, triggering defense responses in infected tissue and signalling intercellular transmission. Upregulation of these genes suggests their involvement in manipulating suppressive activity against the fungal pathogen *R. solani.* We further assessed the expression of five wheat transcripts from the set of highly confident differentially expressed wheat transcripts (with known pathways)-sRNA pairs, which were related to UDPGP, ATP synthase and Ribosomal functions. UDPGP family genes in plants have been associated with a range of functions including pollen mother cell callose deposition, cold-sweetening, tobacco height, chrysolaminaran content, lipid biosynthesis, and carbon allocation [87]. Furthermore, UDGP proteins have been associated with promoting stomatal closure and limiting pathogen entry into leaves. Up-regulated genes like ‘AET1Gv20702500’ and ‘UDPGP’ mediated disease resistance in *Arabidopsis*, supporting immunity [101]. The potential sRNA-UDGP interaction could provide insights into the interplay of rhizosphere microbiome and host defense regulation.

The potential for rhizosphere sRNA mediated crosstalk with host gene expression opens an opportunity for manipulation of host immune responses through management of rhizosphere bacteria. This study provides first experimental support for the potential role of bacterial sRNA-based communication between rhizosphere microorganisms and the host plant with implications for the modulation of disease incidence or severity. Further research could focus on functional validation of *in-planta* gene silencing from sRNAs derived from rhizosphere bacteria, for example through 5 prime RACE, argonaute co-immunoprecipitation, and cleavage assays. Understanding the communication strategies at play within a rhizosphere will facilitate the design of strategies to support the *in situ* development of disease suppressive communities or the engineering of microbiomes that modulate crop gene expression to enhance the sustainability of agriculture food production.

## Acknowledgement

Authors would like to thank CSIRO internal reviewer Dr Ming-Bo Wang for critically reviewing the paper. A special acknowledgement to Dr Scott Rice (Director) MOSH-FSP for his continuous encouragements and feedback in conducting this work.

## Ethics

No animals and humans are involved in this study.

## Availability of data

Raw sequencing datasets will be available in CSIRO Data Access Portal

## References

1. Hakim, S., et al., Rhizosphere engineering with plant growth-promoting microorganisms for agriculture and ecological sustainability. Frontiers in Sustainable Food Systems, 2021. 5: p. 617157.

2. Regmi, R.Y., Qi: Gupta Vadakattu, Advances in understanding microbial communities in the rhizosphere. 2024.

3. Andrews, J.H. and R.F. Harris, The ecology and biogeography of microorganisms on plant surfaces. Annual review of phytopathology, 2000. 38(1): p. 145–180.

4. Mendes, R., et al., Deciphering the rhizosphere microbiome for disease-suppressive bacteria. Science, 2011. 332(6033): p. 1097–1100.

5. Penton, C.R., et al., Fungal Community Structure in Disease Suppressive Soils Assessed by 28S LSU Gene Sequencing. PLOS ONE, 2014. 9(4): p. e93893.

6. Carrión, V.J., et al., Involvement of Burkholderiaceae and sulfurous volatiles in disease-suppressive soils. The ISME journal, 2018. 12(9): p. 2307–2321.

7. Carrión, V.J., et al., Pathogen-induced activation of disease-suppressive functions in the endophytic root microbiome. Science, 2019. 366(6465): p. 606–612.

8. Hayden, H.L., et al., Comparative metatranscriptomics of wheat rhizosphere microbiomes in disease suppressive and non-suppressive soils for Rhizoctonia solani AG8. Frontiers in microbiology, 2018. 9: p. 859.

9. Garrido-Oter, R., et al., Modular traits of the Rhizobiales root microbiota and their evolutionary relationship with symbiotic rhizobia. Cell host & microbe, 2018. 24(1): p. 155–167.e5.

10. Muresu, R., et al., Nodule-associated microbiome diversity in wild populations of Sulla coronaria reveals clues on the relative importance of culturable rhizobial symbionts and co-infecting endophytes. Microbiological research, 2019. 221: p. 10–14.

11. Mushinski, R.M., et al., Nitrogen cycling microbiomes are structured by plant mycorrhizal associations with consequences for nitrogen oxide fluxes in forests. Global change biology, 2021. 27(5): p. 1068–1082.

12. Ikoyi, I., et al., Responses of soil microbiota and nematodes to application of organic and inorganic fertilizers in grassland columns. Biology and Fertility of Soils, 2020. 56: p. 647–662.

13. Xiong, J., et al., Effect of rice (Oryza sativa L.) genotype on yield: Evidence from recruiting spatially consistent rhizosphere microbiome. Soil Biology and Biochemistry, 2021. 161: p. 108395.

14. Wei, N., et al., Genotypic variation in floral volatiles influences floral microbiome more strongly than interactions with herbivores and mycorrhizae in strawberries. Horticulture research, 2022. 9.

15. Chen, Y., et al., Current studies of the effects of drought stress on root exudates and rhizosphere microbiomes of crop plant species. International Journal of Molecular Sciences, 2022. 23(4): p. 2374.

16. Tiziani, R., et al., Drought, heat, and their combination impact the root exudation patterns and rhizosphere microbiome in maize roots. Environmental and Experimental Botany, 2022. 203: p. 105071.

17. Levine, E., et al., Quantitative characteristics of gene regulation by small RNA. PLoS biology, 2007. 5(9): p. e229.

18. Shimoni, Y., et al., Regulation of gene expression by small non-coding RNAs: a quantitative view. Molecular systems biology, 2007. 3(1): p. 138.

19. Mihailovic, M.K., et al., Uncovering Transcriptional Regulators and Targets of sRNAs Using an Integrative Data-Mining Approach: H-NS-Regulated RseX as a Case Study. Front Cell Infect Microbiol, 2021. 11: p. 696533.

20. Regmi, R., et al., Do small RNAs unlock the below ground microbiome-plant interaction mystery? Frontiers in Molecular Biosciences, 2022. 9: p. 1204.

21. Huang, C.-Y., et al., Small RNAs–big players in plant-microbe interactions. Cell host & microbe, 2019. 26(2): p. 173–182.

22. Silvestri, A., et al., In silico analysis of fungal small RNA accumulation reveals putative plant mRNA targets in the symbiosis between an arbuscular mycorrhizal fungus and its host plant. BMC genomics, 2019. 20(1): p. 1–18.

23. Cai, Q., et al., Small RNAs and extracellular vesicles: New mechanisms of cross-species communication and innovative tools for disease control. PLoS pathogens, 2019. 15(12): p. e1008090.

24. Cai, Q., et al., Plants send small RNAs in extracellular vesicles to fungal pathogen to silence virulence genes. Science, 2018. 360(6393): p. 1126–1129.

25. Wang, M., et al., Botrytis small RNA Bc-siR37 suppresses plant defense genes by cross-kingdom RNAi. RNA biology, 2017. 14(4): p. 421–428.

26. Jian, J. and X. Liang, One small RNA of Fusarium graminearum targets and silences CEBiP gene in common wheat. Microorganisms, 2019. 7(10): p. 425.

27. Schaefer, L.K., et al., Cross-kingdom RNAi of pathogen effectors leads to quantitative adult plant resistance in wheat. Frontiers in plant science, 2020. 11: p. 253.

28. Ren, B., et al., Rhizobial tRNA-derived small RNAs are signal molecules regulating plant nodulation. Science, 2019. 365(6456): p. 919–922.

29. Wen, H.-G., et al., Microbe-induced gene silencing boosts crop protection against soil-borne fungal pathogens. Nature Plants, 2023. 9(9): p. 1409–1418.

30. Middleton, H., et al., Rhizospheric plant–microbe interactions: miRNAs as a key mediator. Trends in plant science, 2021. 26(2): p. 132–141.

31. Donn, S., et al., Rhizosphere microbial communities associated with Rhizoctonia damage at the field and disease patch scale. Applied soil ecology, 2014. 78: p. 37–47.

32. Davey, R.S., et al., Potential for suppression of Rhizoctonia root rot is influenced by nutrient (N and P) and carbon inputs in a highly calcareous coarse-textured topsoil. Soil Research, 2021. 59(4): p. 329–345.

33. Murray, G. and J. Brennan, Estimating disease losses to the Australian barley industry. Australasian Plant Pathology, 2010. 39(1): p. 85–96.

34. Gupta, V.V., Beneficial microorganisms for sustainable agriculture. Microbiology Australia, 2012. 33(3): p. 113–115.

35. Liu, H., et al., Inner plant values: diversity, colonization and benefits from endophytic bacteria. Frontiers in microbiology, 2017. 8: p. 2552.

36. Andrews, S., FastQC: a quality control tool for high throughput sequence data. 2010, Babraham Bioinformatics, Babraham Institute, Cambridge, United Kingdom.

37. Bushnell, B., BBMap: a fast, accurate, splice-aware aligner. 2014, Lawrence Berkeley National Lab.(LBNL), Berkeley, CA (United States).

38. Quast, C., et al., The SILVA ribosomal RNA gene database project: improved data processing and web-based tools. Nucleic acids research, 2012. 41(D1): p. D590–D596.

39. Axtell, M.J., ShortStack: comprehensive annotation and quantification of small RNA genes. Rna, 2013. 19(6): p. 740–751.

40. Grabherr, M.G., et al., Full-length transcriptome assembly from RNA-Seq data without a reference genome. Nature biotechnology, 2011. 29(7): p. 644–652.

41. Li, D., et al., MEGAHIT: an ultra-fast single-node solution for large and complex metagenomics assembly via succinct de Bruijn graph. Bioinformatics, 2015. 31(10): p. 1674–1676.

42. Wu, Y.-W., B.A. Simmons, and S.W. Singer, MaxBin 2.0: an automated binning algorithm to recover genomes from multiple metagenomic datasets. Bioinformatics, 2016. 32(4): p. 605–607.

43. Nawrocki, E.P. and S.R. Eddy, Infernal 1.1: 100-fold faster RNA homology searches. Bioinformatics, 2013. 29(22): p. 2933–2935.

44. Regmi, R., et al., Identification of B. napus small RNAs responsive to infection by a necrotrophic pathogen. BMC Plant Biol, 2021. 21(1): p. 366.

45. Langmead, B. and S.L. Salzberg, Fast gapped-read alignment with Bowtie 2. Nature methods, 2012. 9(4): p. 357–359.

46. Wood, D.E., J. Lu, and B. Langmead, Improved metagenomic analysis with Kraken 2. Genome Biology, 2019. 20(1): p. 257.

47. Breitwieser, F.P. and S.L. Salzberg, Pavian: interactive analysis of metagenomics data for microbiome studies and pathogen identification. Bioinformatics, 2020. 36(4): p. 1303–1304.

48. Dai, X., Z. Zhuang, and P.X. Zhao, psRNATarget: a plant small RNA target analysis server (2017 release). Nucleic acids research, 2018. 46(W1): p. W49–W54.

49. Raden, M., et al., Freiburg RNA tools: a central online resource for RNA-focused research and teaching. Nucleic acids research, 2018. 46(W1): p. W25–W29.

50. Sherman, B.T., et al., DAVID: a web server for functional enrichment analysis and functional annotation of gene lists (2021 update). Nucleic acids research, 2022. 50(W1): p. W216–W221.

51. Wickham, H., ggplot2. Wiley interdisciplinary reviews: computational statistics, 2011. 3(2): p. 180–185.

52. Huerta-Cepas, J., et al., eggNOG 5.0: a hierarchical, functionally and phylogenetically annotated orthology resource based on 5090 organisms and 2502 viruses. Nucleic acids research, 2019. 47(D1): p. D309–D314.

53. Bolser, D., et al., Ensembl plants: integrating tools for visualizing, mining, and analyzing plant genomics data. Plant bioinformatics: Methods and protocols, 2016: p. 115–140.

54. Derbyshire, M., et al., Small RNAs from the plant pathogenic fungus Sclerotinia sclerotiorum highlight host candidate genes associated with quantitative disease resistance. Mol Plant Pathol, 2019. 20(9): p. 1279–1297.

55. Regmi, R., et al., Genome-wide identification of Sclerotinia sclerotiorum small RNAs and their endogenous targets. BMC Genomics, 2023. 24(1): p. 582.

56. de Haro, L.A., et al., Mal de Río Cuarto virus infection triggers the production of distinctive viral-derived siRNA profiles in wheat and its planthopper vector. Frontiers in plant science, 2017. 8: p. 766.

57. Ma, X., J. Wiedmer, and J. Palma-Guerrero, Small RNA bidirectional crosstalk during the interaction between wheat and Zymoseptoria tritici. Frontiers in plant science, 2020. 10: p. 1669.

58. Gottesman, S. and G. Storz, Bacterial small RNA regulators: versatile roles and rapidly evolving variations. Cold Spring Harbor perspectives in biology, 2011. 3(12): p. a003798.

59. Huen, A., J. Bally, and P. Smith, Identification and characterisation of microRNAs and their target genes in phosphate-starved Nicotiana benthamiana by small RNA deep sequencing and 5’RACE analysis. BMC Genomics, 2018. 19(1): p. 940.

60. Thomas, P.D., The gene ontology and the meaning of biological function. The gene ontology handbook, 2017: p. 15–24.

61. Li, B. and C.N. Dewey, RSEM: accurate transcript quantification from RNA-Seq data with or without a reference genome. BMC Bioinformatics, 2011. 12(1): p. 323.

62. Kalvari, I., et al., Rfam 14: expanded coverage of metagenomic, viral and microRNA families. Nucleic Acids Research, 2021. 49(D1): p. D192–D200.

63. Sass, A., S. Kiekens, and T. Coenye, Identification of small RNAs abundant in Burkholderia cenocepacia biofilms reveal putative regulators with a potential role in carbon and iron metabolism. Scientific reports, 2017. 7(1): p. 15665.

64. Greenlee, E.B., et al., Challenges of ligand identification for the second wave of orphan riboswitch candidates. RNA biology, 2018. 15(3): p. 377–390.

65. Bloch, S., et al., Small and Smaller-sRNAs and MicroRNAs in the Regulation of Toxin Gene Expression in Prokaryotic Cells: A Mini-Review. Toxins (Basel), 2017. 9(6).

66. Jin, L., et al., The molecular mechanism of dsRNA processing by a bacterial Dicer. Nucleic Acids Res, 2019. 47(9): p. 4707–4720.

67. Vandivier, L.E., et al., The Conservation and Function of RNA Secondary Structure in Plants. Annu Rev Plant Biol, 2016. 67: p. 463–88.

68. McKellar, S.W., et al., RNase III CLASH in MRSA uncovers sRNA regulatory networks coupling metabolism to toxin expression. Nature Communications, 2022. 13(1): p. 3560.

69. Schlatter, D., et al., Disease Suppressive Soils: New Insights from the Soil Microbiome. Phytopathology®, 2017. 107(11): p. 1284–1297.

70. Mendes, L.W., et al., Influence of resistance breeding in common bean on rhizosphere microbiome composition and function. The ISME Journal, 2018. 12(1): p. 212–224.

71. Bousios, A., B.S. Gaut, and N. Darzentas, Considerations and complications of mapping small RNA high-throughput data to transposable elements. Mobile DNA, 2017. 8: p. 1–13.

72. Neeb, Zachary T. and Alan M. Zahler, An Expanding World of Small RNAs. Developmental Cell, 2014. 28(2): p. 111–112.

73. Jia, J., et al., The 25–26 nt small RNAs in Phytophthora parasitica are associated with efficient silencing of homologous endogenous genes. Frontiers in microbiology, 2017. 8: p. 773.

74. Sun, Z., et al., tRNA-derived fragments from wheat are potentially involved in susceptibility to Fusarium head blight. BMC Plant Biology, 2022. 22(1): p. 1–17.

75. Mi, S., et al., Sorting of small RNAs into Arabidopsis argonaute complexes is directed by the 5′ terminal nucleotide. Cell, 2008. 133(1): p. 116–127.

76. Sánchez-Correa, M.d.S., et al., Argonaute5 and its associated small RNAs modulate the transcriptional response during the rhizobia-Phaseolus vulgaris symbiosis. Frontiers in Plant Science, 2022. 13: p. 1034419.

77. Liu, Y.-F., et al., Genome-wide identification and evolutionary analysis of Argonaute genes in hexaploid bread wheat. BioMed Research International, 2021. 2021: p. 1–9.

78. Sibisi, P. and E. Venter, Wheat argonaute 5 functions in aphid–plant interaction. Frontiers in Plant Science, 2020. 11: p. 641.

79. He, B., et al., Botrytis cinerea small RNAs are associated with tomato AGO1 and silence tomato defense-related target genes supporting cross-kingdom RNAi. bioRxiv, 2023: p. 2022.12.30.522274.

80. Collins, C.M. and S.E. D’Orazio, Bacterial ureases: structure, regulation of expression and role in pathogenesis. Molecular microbiology, 1993. 9(5): p. 907–913.

81. Pantigoso, H.A., D. Newberger, and J.M. Vivanco, The rhizosphere microbiome: Plant-microbial interactions for resource acquisition. J Appl Microbiol, 2022. 133(5): p. 2864–2876.

82. Veaudor, T., C. Cassier-Chauvat, and F. Chauvat, Genomics of Urea Transport and Catabolism in Cyanobacteria: Biotechnological Implications. Frontiers in Microbiology, 2019. 10.

83. Wu, Y., et al., Suppression of host plant defense by bacterial small RNAs packaged in outer membrane vesicles. Plant Communications, 2024. 5(4): p. 100817.

84. Seth, T., et al., The intricate role of lipids in orchestrating plant defense responses. Plant Science, 2024. 338: p. 111904.

85. Jagodzik, P., et al., Mitogen-Activated Protein Kinase Cascades in Plant Hormone Signaling. Frontiers in Plant Science, 2018. 9.

86. Kang, J., et al., Plant ABC transporters. The Arabidopsis book/American Society of Plant Biologists, 2011. 9.

87. Hung, C.-Y., et al., Phosphoinositide-signaling is one component of a robust plant defense response. Frontiers in Plant Science, 2014. 5: p. 267.

88. Berens, M.L., et al., Evolution of hormone signaling networks in plant defense. Annual review of phytopathology, 2017. 55: p. 401–425.

89. Qasim, M.U., et al., Overlapping pathways involved in resistance against Sclerotinia stem rot in Brassica napus revealed through transcriptomic and metabolomic profiling. Plant Growth Regulation, 2023: p. 1–16.

90. Wei, L., et al., Integrated mRNA and miRNA transcriptome analysis of grape in responses to salt stress. Front Plant Sci, 2023. 14: p. 1173857.

91. Campos, C., et al., Transcriptome Analysis of Wheat Roots Reveals a Differential Regulation of Stress Responses Related to Arbuscular Mycorrhizal Fungi and Soil Disturbance. Biology (Basel), 2019. 8(4).

92. Wang, J., et al., Transcriptome analysis in roots and leaves of wheat seedlings in response to low-phosphorus stress. Scientific Reports, 2019. 9(1): p. 19802.

93. Xi, W., et al., Transcriptome Analysis of Roots from Wheat (Triticum aestivum L.) Varieties in Response to Drought Stress. International Journal of Molecular Sciences, 2023. 24(8): p. 7245.

94. Jing, Y., et al., Plant elicitor peptide induces endocytosis of plasma membrane proteins in Arabidopsis. Frontiers in Plant Science, 2023. 14.

95. Wilson, D.F., Oxidative phosphorylation: regulation and role in cellular and tissue metabolism. The Journal of Physiology, 2017. 595(23): p. 7023–7038.

96. Kimberlin, A., R.E. Holtsclaw, and A.J. Koo, Differential regulation of the ribosomal association of mRNA transcripts in an Arabidopsis mutant defective in jasmonate-dependent wound response. Frontiers in plant science, 2021. 12: p. 637959.

97. Huang, X.X., et al., The novel pathogen-responsive glycosyltransferase UGT73C7 mediates the redirection of phenylpropanoid metabolism and promotes SNC1- dependent Arabidopsis immunity. The Plant Journal, 2021. 107(1): p. 149–165.

98. Hafez, E. and M. Moustafa, Differential expression of P19, cytochrome oxidase and ALY-family genes in resistant and susceptible tomato cultivars (Solanum lycopersicum) inoculated with tomato bushy stunt virus. Genomics and Applied Biology, 2011. 2(2).

99. Palmer, A.J., A. Baker, and S.P. Muench, The varied functions of aluminium-activated malate transporters–much more than aluminium resistance. Biochemical Society Transactions, 2016. 44(3): p. 856–862.

100. Ramesh, S.A., et al., Aluminum-activated malate transporters can facilitate GABA transport. The Plant Cell, 2018. 30(5): p. 1147–1164.

101. Tang, D., G. Wang, and J.-M. Zhou, Receptor Kinases in Plant-Pathogen Interactions: More Than Pattern Recognition. The Plant Cell, 2017. 29(4): p. 618–637.

